# Experience-dependent structural plasticity at pre- and postsynaptic sites of layer 2/3 cells in developing visual cortex

**DOI:** 10.1101/743690

**Authors:** Yujiao Jennifer Sun, J. Sebastian Espinosa, Mahmood S. Hoseini, Michael P. Stryker

## Abstract

The developing brain can respond quickly to altered sensory experience by circuit reorganization. During a critical period in early life, neurons in the primary visual cortex rapidly lose responsiveness to an occluded eye and come to respond better to the open eye. While physiological and some of the molecular mechanisms of this process have been characterized, its structural basis, except for the well-known changes in the thalamocortical projection, remains obscure. To elucidate the relationship between synaptic remodeling and functional changes during this experience-dependent process, we used 2-photon microscopy to image synaptic structures of sparsely labeled layer 2/3 neurons in the binocular zone of mouse primary visual cortex. Anatomical changes at presynaptic and postsynaptic sites in mice undergoing monocular visual deprivation (MD) were compared to those in control mice with normal visual experience. We found that postsynaptic spines remodeled quickly in response to MD, with neurons more strongly dominated by the deprived eye losing more spines. These postsynaptic changes parallel changes in visual responses during MD and their recovery after restoration of binocular vision. In control animals with normal visual experience, the formation of presynaptic boutons increased during the critical period and then declined. MD affected bouton formation, but with a delay, blocking it after 3 days. These findings reveal intracortical anatomical changes in cellular layers of the cortex that can account for rapid activity-dependent plasticity.

**Significance statement:** The operation of the cortex depends on the connections among its neurons. Taking advantage of molecular and genetic tools to label major proteins of the presynaptic and postsynaptic densities, we studied how connections of layer 2/3 excitatory neurons in mouse visual cortex were changed by monocular visual deprivation during the critical period, which causes amblyopia. The deprivation induced rapid remodeling of postsynaptic spines and impaired bouton formation. Structural measurement followed by calcium imaging demonstrated a strong correlation between changes in postsynaptic structures and functional responses in individual neurons after monocular deprivation. These findings suggest that anatomical changes at postsynaptic sites serve as a substrate for experience-dependent plasticity in the developing visual cortex.

## Introduction

Sensory experience during early life plays an instructive role in the formation and maturation of neural circuits (1–4). Abnormal experience within early critical periods have life-long effects (5–7). In the visual system, monocular visual deprivation (MD) within a critical period causes drastic changes in visual cortical responses in many species (8–12) that are highly stereotyped and reproducible. In the mouse, the effects of MD during the critical period take place in temporally and mechanistically distinct stages: a Hebbian-like process with decreased response to the deprived eye in the first 2 to 3 days (13), followed by a delayed increase in response to the open eye governed by both Hebbian and homeostatic rules (14–17). After reopening the deprived eye within the critical period, recovery of responses to normal levels occurs within 2 days (15, 18, 19).

Structural reorganization at various anatomical sites has also been described in association with the overall loss of response to the deprived eye. At thalamorecipient layer 4, remodeling of thalamic axonal arbors takes place over days to weeks (cat (20–23), mouse (24)). Physiological changes beyond the input layer of cat visual cortex are much more rapid (25), and the reorganization of intracortical connections (26) in the extragranular layers has a similar time course to the rapid changes in physiological responses. These findings support the idea that synapses beyond layer 4 are the loci for rapid remodeling (25). In both the adult (27–32) and developing (33) visual cortex of the mouse, altered experience increases turnover of excitatory and inhibitory synapses in the superficial layers. Population studies of spines in layer 1 *in vivo* and spines in deeper layers *in vitro* have shown that prolonged dark-rearing (34, 35) or MD (36, 37) decreases spine numbers in the visual cortex. However, the extent to which rewiring of upper-layer connections accounts for the functional changes in individual neurons remains unclear.

To answer this question, we studied synaptic remodeling longitudinally in individual neurons of the binocular zone of mouse visual cortex (V1) over the stages of ocular dominance plasticity during critical period. We used 2-photon imaging to track the anatomical rearrangement of presynaptic boutons and postsynaptic spines within layer 2/3 excitatory neurons. Comparing the changes in control mice receiving normal visual input to those in experimental mice undergoing monocular deprivation and binocular recovery, we found that postsynaptic spines respond quickly to the altered visual experience and that this process is strongly correlated with the functional responses of individual cells. Presynaptic boutons also respond to altered visual experience but with a longer latency. The developmental increase in bouton formation was disrupted only after 3 days of MD.

## Results

### Labeling synaptic structures for longitudinal imaging

To study experience-dependent structural plasticity in the developing mouse visual cortex, we sparsely labeled layer 2/3 cortical excitatory neurons for longitudinal imaging. We designed Cre-recombinase-dependent plasmids with GFP fused to proteins that are abundant presynaptically (synaptophysin, SYP) or postsynaptically (the 95kd postsynaptic density protein, PSD95) to mark pre- and postsynaptic structures in individual neurons. To allow long-term visualization of these structures in layer 2/3 excitatory neurons *in vivo*, either PSD95-GFP or SYP-GFP plasmids were delivered together with Cre-recombinase into E16 embryos of Ai14 mice, which conditionally express tdTomato (Fig 1A). A cranial window with a cover glass was implanted over the primary visual cortex one week before the structural imaging, and intrinsic signal imaging was first carried out to identify the binocular zone (Fig 1B). Around the peak of the critical period, near postnatal day 28 (P28) (10), we used 2-photon microscopy to track longitudinal changes in individual spines or boutons of cortical neurons in animals experiencing normal vision or MD.

**Figure 1.**
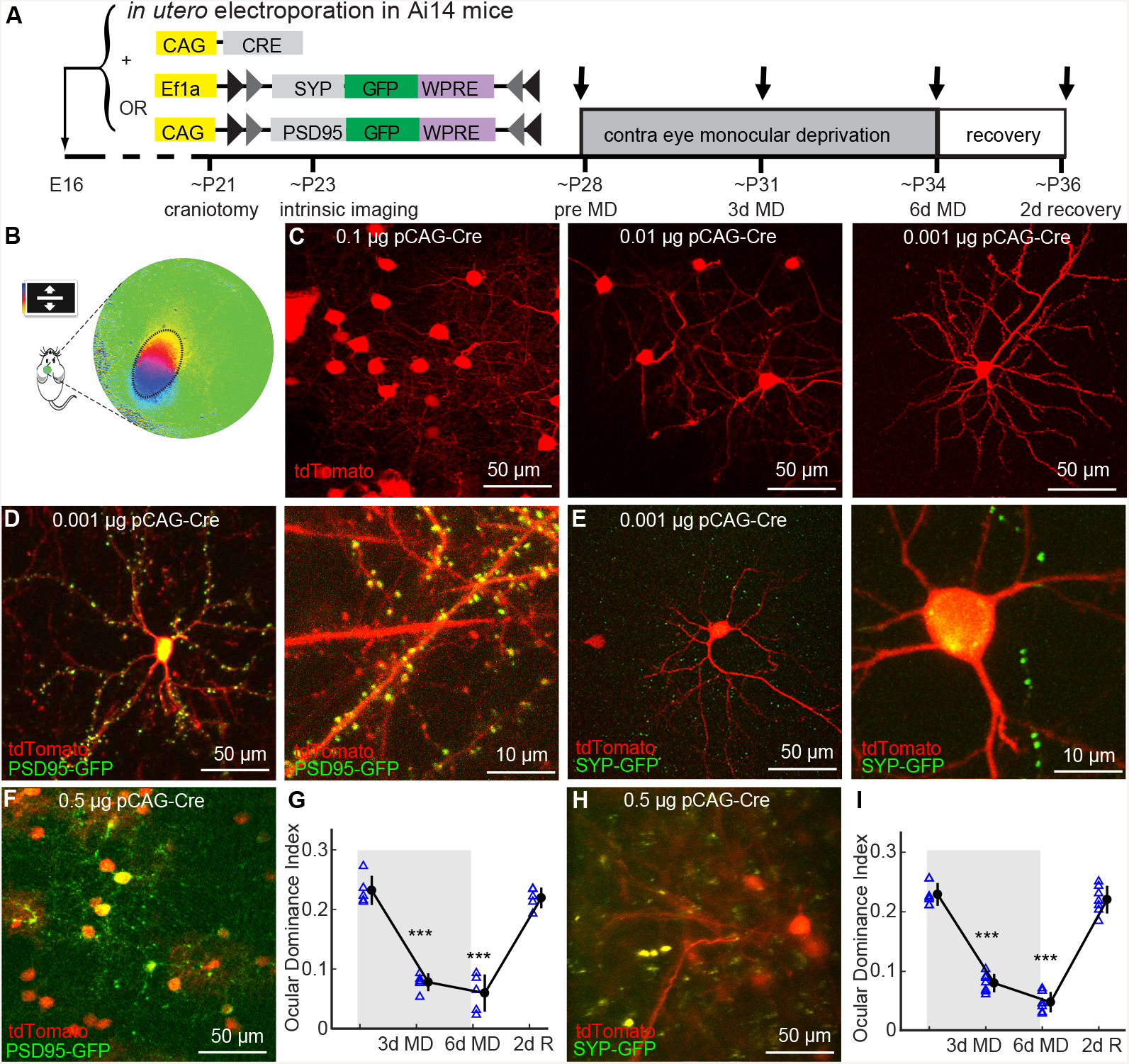
Labeling of synaptic markers does not affect functional changes during MD. A, Timeline of experiments: at embryonic day 16, plasmids (pCAG-Cre and double floxed markers, either SYP-GFP or PSD95-GFP) were delivered into left ventricle in Ai14 embryos. After craniotomy around postnatal day (P) 21, cortex was first screened for sparse neuronal labeling. Intrinsic imaging then was used to determine the exact location of the binocular zone of primary visual cortex. Monocular depriva- (MD) of experimental animals was performed by suture of contralateral eyelid around P25 and lasted for 6 days. Structural imaging was performed at times indicated by arrows. B, Example of intrinsic signal imaging to identify binocular zone of visual cortex. C, Different dosages of pCAG-Cre plasmid delivered in Ai14 animals through *in utero* electroporation were tested for sparse labeling; 0.001μg pCAG-Cre was chosen for structural imaging. D, of pCAG-Cre and double floxed postsynaptic marker PSD95-GFP labeled cell bodies and neurite structure in red channel and postsynaptic structures in green channel. E, Similar to D but for presynaptic marker SYP-GFP. F, Overexpression of Cre (0.5 μg/μl) and PSD95-GFP (0.75 μg/μl) in Ai14 mice resulted high density labeling of cells and postsynaptic markers PSD95-GFP. G, Intrinsic signal imaging demonstrated normal ODI shift with PSD95-GFP overexpression, each represents a single animal. HI, Similar to FG but for presynaptic marker, overexpression of Cre and SYP-GFP in Ai14 animals resulted normal ODI shift. Error bar represents mean ± S.D. ***p < 0.001 compared with the baseline (day 0) response (one-way ANOVA followed by multiple comparisons with Tukey’s method).

To achieve the sparse labeling, a range of pCAG-Cre concentrations was delivered to Ai14 mouse embryos in which all neurons contain floxed tdTomato (Fig 1C). A low concentration of the plasmid expressing Cre-recombinase (0.001μg) yielded sparse labeling of individual neurons and was used for longitudinal structural imaging. The sparsely labeled layer 2/3 neurons were marked with red tdTomato fluorescence in their cell bodies, dendrites, and axons. Mixing the low concentration of recombinase plasmid with a higher concentration of the plasmid encoding either SYP-GFP or PSD95-GFP ensured that pre-or postsynaptic structures were also labeled with GFP in all the red neurons. Neurons with PSD95-GFP labeling express bright fluorescent puncta within individual spines viewed using 2-photon imaging (Fig 1D). In neurons with SYP-GFP labeling, presynaptic boutons are evident as isolated puncta on the axons marked with red fluorescence (Fig 1E).

### GFP-tagged synaptic markers do not impair ocular dominance plasticity

A control experiment was carried out to confirm that the GFP-labeled synaptic markers did not alter the normal functional plasticity of mouse visual cortex. We labeled layer 2/3 neurons densely rather than sparsely with PSD95-GFP or SYP-GFP and measured ocular dominance plasticity using intrinsic signal imaging (Fig 1B). Ocular dominance plasticity during MD and recovery phases was unaffected even by the 500-fold higher labeling of postsynaptic (Fig 1G) or presynaptic (Fig 1I) sites than was used in the experimental conditions (15, 16).

### Image processing pipeline and semi-automated puncta detection

In many cases, up to 9 overlapping volume image stacks were acquired in the same imaging session to capture a labeled neuron and its synaptic structures. The image stacks were first stitched together to make a larger single image using rigid translation based on the red fluorescence channel (Fig 2B). Composite images acquired over 3 additional imaging sessions at 2-3 day intervals were then aligned to the baseline images by non-rigid warping of the red channel using procedures in the Computational Morphology Toolkit (CMTK) (38). Images acquired even 8 days apart were aligned by this procedure (Fig 2C).

**Figure 2.**
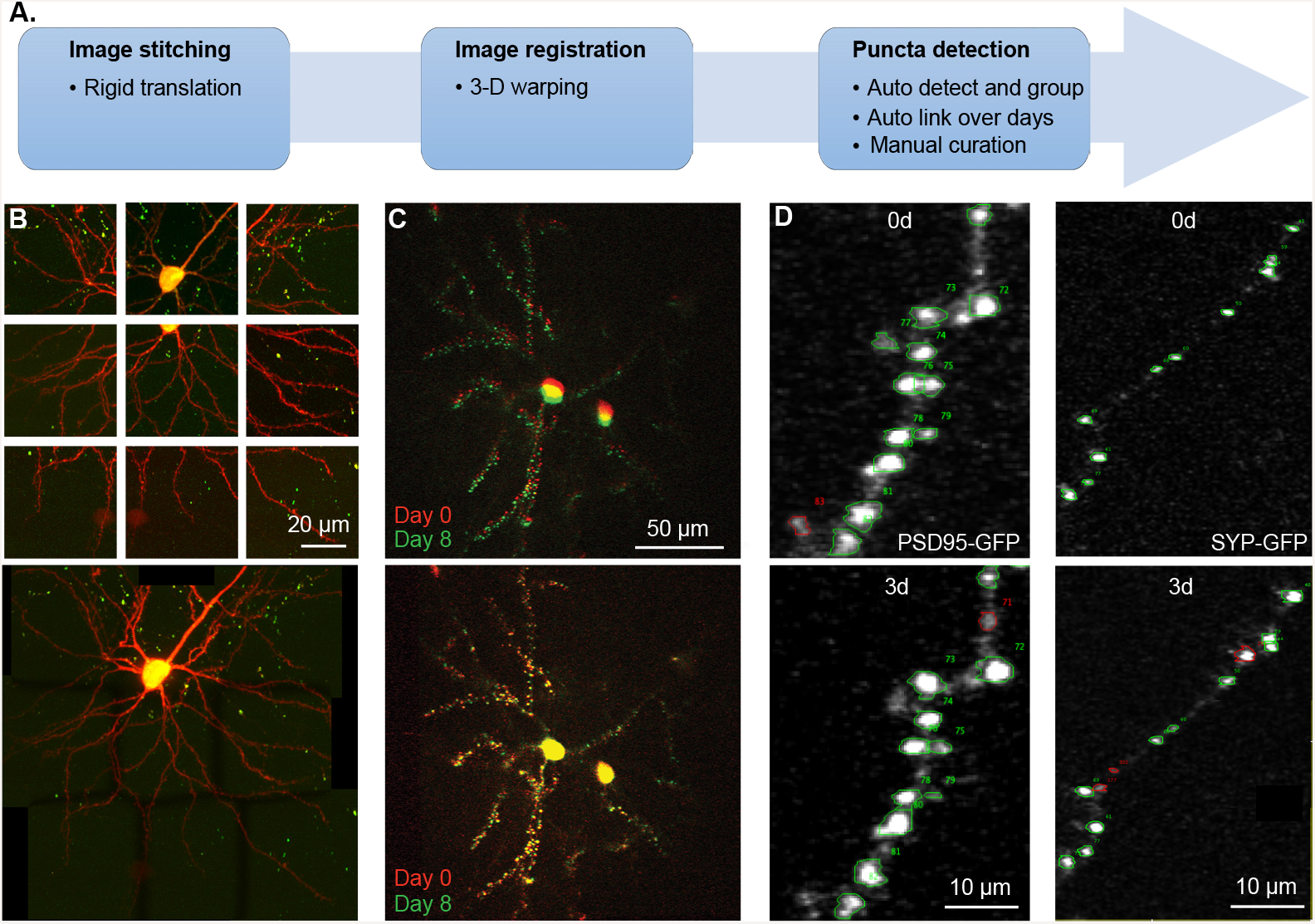
Image processing pipeline and semi-automated puncta detection. A, Image processing pipeline consists of three steps: image stitching, image registration, and semi-automatic puncta detection. B, Nine high-resolution images were taken covering the region of interest (ROI) in a layer 2/3 excitatory neuron (top panel), and automated 3D stitching using rigid translation was applied to obtain a single ROI (bottom panel). C, tdTomato channel images taken over days false-colored in red and green and superimposed (top panel) were aligned using 3D warping (non-rigid registration) to correct the misalignment caused by imaging acquisition as well as tissue growth (bottom panel). D, Automated detection and grouping of 3D puncta were performed by comparing ROIs over different days. Puncta detected over different days were automatically linked based on their spatial locations, followed by manual inspection to verify and make necessary correction. Examples of PSD95-GFP and SYP-GFP are shown, with green circle denoting the linked puncta and red circle denoting the unlinked ones.

To measure large members of pre- and postsynaptic puncta, a custom ImageJ plugin was written for semi-automated puncta detection and measurement (Fig 2D, see Methods). Because GFP expression is confined to synaptic sites, but was not present at detectable levels in our images of dendrites and axons, it appears as puncta in the green channel that are reliably detected using our algorithm. The detected puncta were then linked over days based on criteria optimized either for post- or presynaptic markers (Fig 2D). This high-throughput procedure, which allowed us to process the images to a reproducible and consistent standard, was in all cases followed by manual inspection to verify and make corrections when necessary.

### MD causes bimodal changes in postsynaptic densities

To examine the anatomical changes underlying the different stages of ocular dominance plasticity, we tracked the turn-over of postsynaptic structures in layer 2/3 neurons through the critical period (~P28 – P36), at intervals of 2-3 days (15, 16). We studied 4 control mice with normal visual experience for 8 days and 4 MD mice in which the contralateral eyelid was sutured closed for 6 days then reopened for the following 2 days.

Previous physiological experiments showed that this procedure produces a progressive and robust ocular dominance shift that is reversed by the 2 days of binocular vision (15, 16). In the MD mice, the first 3 days of MD at the peak of critical period caused a drastic change in the spine densities (Fig 3A): many spines disappeared (marked by blue arrow) and few new spines formed (red arrow). Spine elimination was significantly increased (15.6±1.8% in 3d MD vs. 7.9±0.6% in 3d control, p < 0.0001, n=20 dendrites in 4 experimental animals, n=19 dendrites in 4 control animals, unpaired Welch’s t-test) but there was only a modest, insignificant change in spine formation (6.9±1.2% in 3d MD vs 4.6±0.5% in 3d control, p = 0.066). As a result, 3 days of MD caused a net loss of spines that was significantly larger than control (−8.7±2.4% in 3d MD vs −3.3±0.9% in 3d control, p = 0.035). In contrast, spine elimination and formation were balanced over this period in the control animals (Fig 3 D-F).

A delayed but significant increase in spine formation was evident by 6 days of MD (12.2±2.0% in 6d MD vs 5.8±0.9% in 6d control, p = 0.0035), in addition to the spine elimination (12.1±1.7% in 6d MD vs 8.0±0.8% in 6d control, p = 0.029). Consistent with the different direction of changes in the responses of neurons dominated by the deprived and nondeprived eyes (10), the net spine changes at 6 days of MD appear bimodal, with some dendritic branches losing many spines and others growing many (Fig 3D, see discussion). This bimodality (or multi-modality) was evident in that the different spine changes among the dendrites could no longer be fit by a normal distribution (p = 0.0123 in 6d MD, p= 0.26 in 6d control, Lilliefors test for normal distribution). The diversity of changes of spines among the dendrites was reversed by 2 days of binocular vision, resulting in a rate of spine elimination and formation similar to control animals (Fig 3B-D, elimination: 7.6±1.4% in 2d recovery vs 6.1±0.9% at the corresponding time in control mice; formation: 10.3±1.0% in 2d recovery vs 9.3±0.9% in control; net change: −3.1 ±2.0% in 2d recovery vs −3.2±1.4% in control).

**Figure 3.**
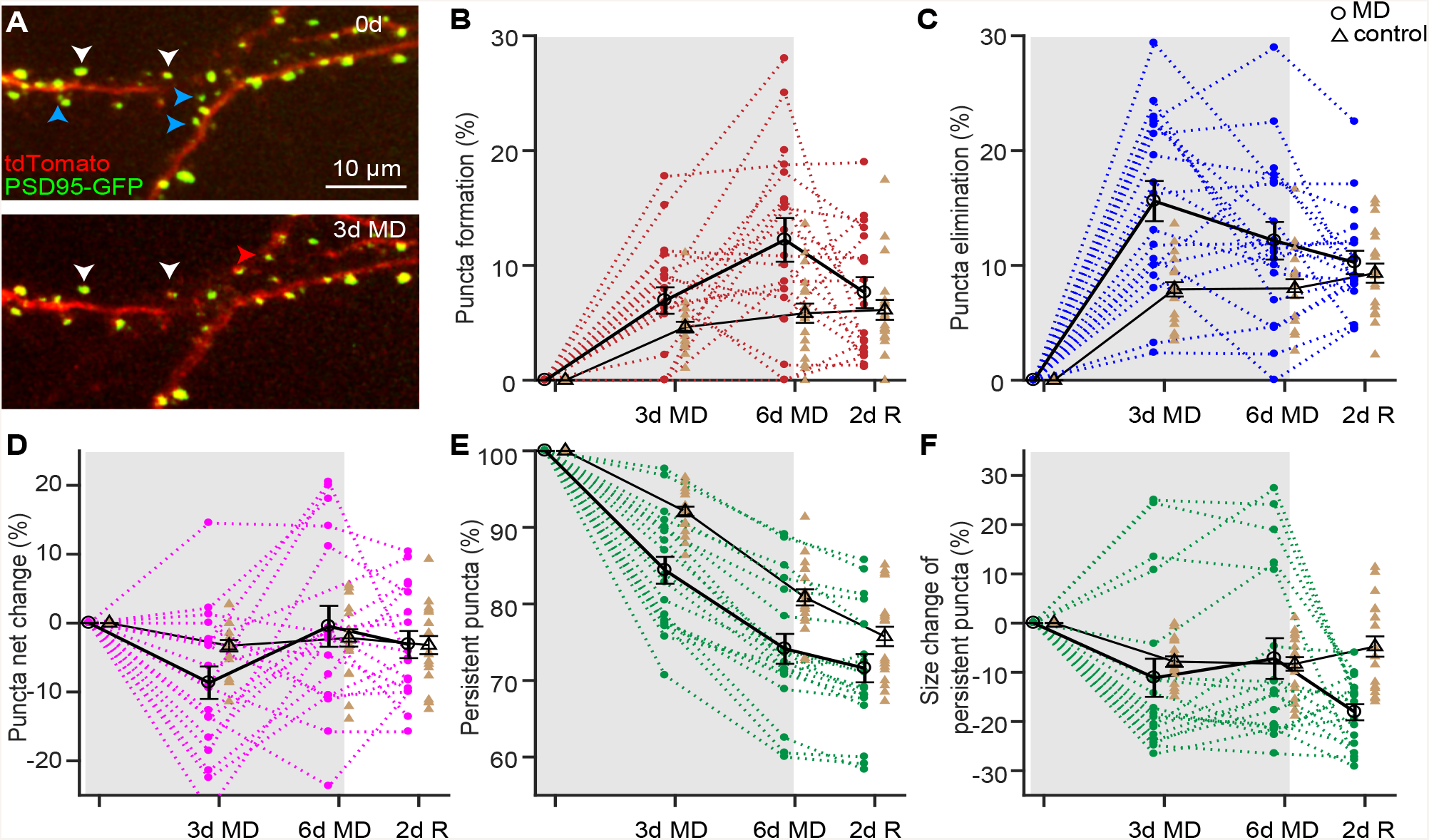
Changes in postsynaptic densities of layer 2/3 neurons. A, Postsynaptic marker PSD95-GFP co-expressed with tdTomato in a layer 2/3excitatory neuron. White arrows mark persistent boutons, blue arrows mark lost bouton, and a red arrow marks new bouton. B-D, Percentage change in dendritic spine formation, elimination, and total net change over 8 days in the binocular zone. E, Fraction of persistent PSD95-GFP puncta throughout the period of MD and recovery. F, Percentage change in size of persistent PSD95-GFP puncta. Each point marks the average value for the 18-54 spines on a single dendrite. Successive measurements of each dendrite in experimental animals connected by dotted lines. Average values for the 20 dendrites in 4 experimental animals indicated by open circles and heavy black line (627 puncta total). Control data plotted in light brown; average values for the 19 dendrites in 4 control animals indicated by open triangles connected by light black lines (711 puncta total).

Over the 8 days of imaging, there were significantly fewer persistent puncta in MD mice than in control mice (84.4±1.8% in 3d MD vs 92.1±0.6% in 3d control, p = 0.0045; 74.1±2.0% in 6d MD vs 80.9±1.1% in 6d control, p = 0.0223, 2-way ANOVA corrected for multiple comparisons), indicating the altered visual experience destabilized the existing puncta (Fig 3E). After 2 days of normal vision, this difference became insignificant (71.6±1.9% in 2d recovery vs. 75.8±1.3% in 8d control, p = 0.3526). The change in size of the persistent puncta (Fig 3F) in the deprived mice also appears bimodal, with some puncta growing bigger while others became smaller (p = 0.0061 in 3d MD, p = 0.016 in 6d MD, p = 0.5 in 3d control, p= 0.49 for 6d control, Lilliefors test for normal distribution). Interestingly, restoration of normal vision for 2 days was sufficient for the enlarged puncta to revert to normal size, but not for the reduced puncta to recover, resulting in a net decrease in puncta size (−18.1±1.7% in 2d recovery vs. −4.8±2.1% in 8d control, p = 0.0058, 2-way ANOVA corrected for multiple comparisons).

### Structural changes of postsynaptic densities correlate with their functional responses

To elucidate the relationship between the anatomical changes and function during ocular dominance plasticity, we measured both in the same layer 2/3 excitatory neurons in 4 additional animals. We first examined the changes in dendritic spines produced by MD by imaging neurons that co-expressed PSD95-GFP and tdTomato before and after 3 days of deprivation. After tracking the spine changes, we performed 2-photon calcium imaging to measure the functional responses in the same neurons. The calcium indicator Oregon Green Bapta-AM (OGB1) was applied to the binocular zone of V1 after removing the cover glass from the cranial window. Full-field drifting gratings were shown successively to the two eyes in order to elicit eye-specific visual responses. The responses at the preferred orientation for each cell were used to calculate the neuron’s ocular dominance index (ODI) (Fig 4A). Such structural and functional measurements revealed a strong correlation between the anatomical changes in spines and the eye-specific responses: dendrites of neurons more strongly dominated by the deprived eye (higher ODI) lost more PSD95-GFP puncta, while those of neurons in which responses to the two eyes were more nearly equal (lower ODI) lost fewer or gained puncta (Fig 4B).

**Figure 4.**
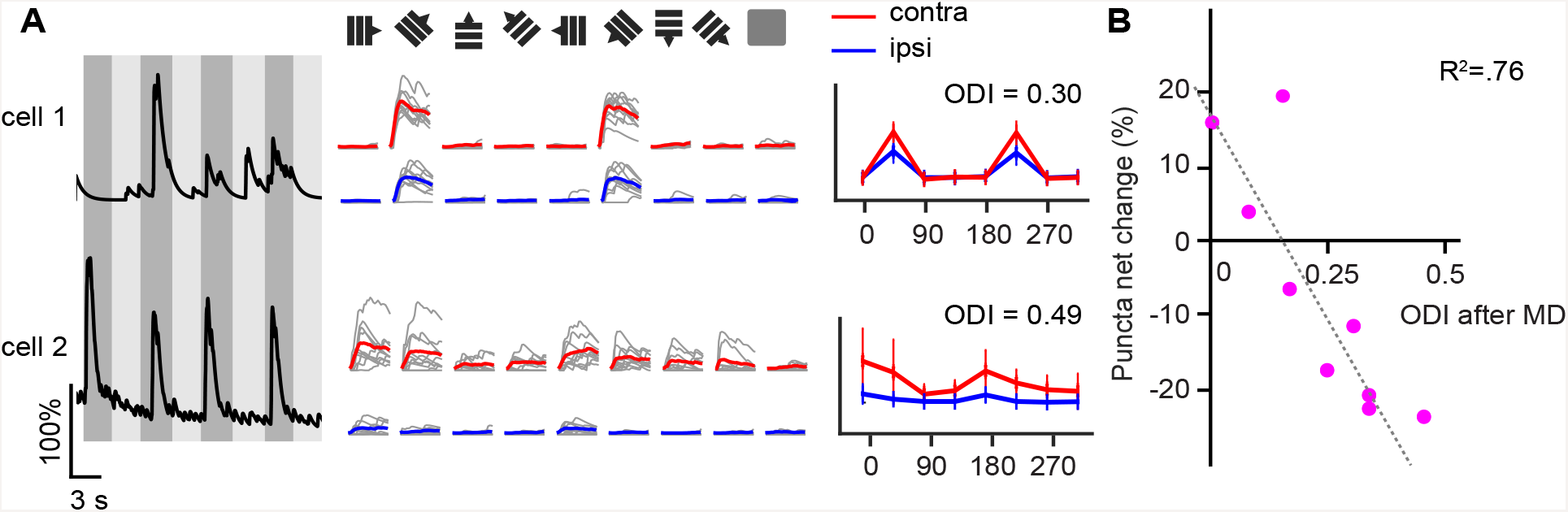
Relationship of ocular dominance in single layer 2/3 neurons to postsynaptic structural changes during MD. A, Functional imaging after structural imaging following 3d MD in a separate group of 4 animals. Full-field drifting gratings of 8 different orientations presented to left and right eye to evoke eye-specific responses. Averaged response at the preferred orientation was used to calculate the ocular dominance index in each neuron B, The loss/gain of postsynaptic density marker PSD95-GFP in each neuron was strongly correlated with the ocular dominance index (r2 = 0.76). 9 neurons with 143-1123 puncta per neuron.

### MD reduces formation of presynaptic boutons

We examined the turnover of presynaptic boutons within layer 2/3 in excitatory neurons of that layer that co-expressed SYP-GFP and tdTomato (Fig 5A). In control mice, there was a steady, significant net increase in the number of presynaptic boutons until the peak of critical period, followed by a return to baseline at the end of the critical period (Fig 5B-D, 6.3 ± 2.1% in 3d control, 11.2 ± 3.1% in 6d control, −3.5 ± 2.8% in 8d control, n = 24 branches in 3 control animals, p = 0.011 for 3d vs. baseline, p < 0.001 for 6d vs. baseline, p = 0.60 for 8d vs. baseline, ANOVA corrected for multiple comparisons).

**Figure 5.**
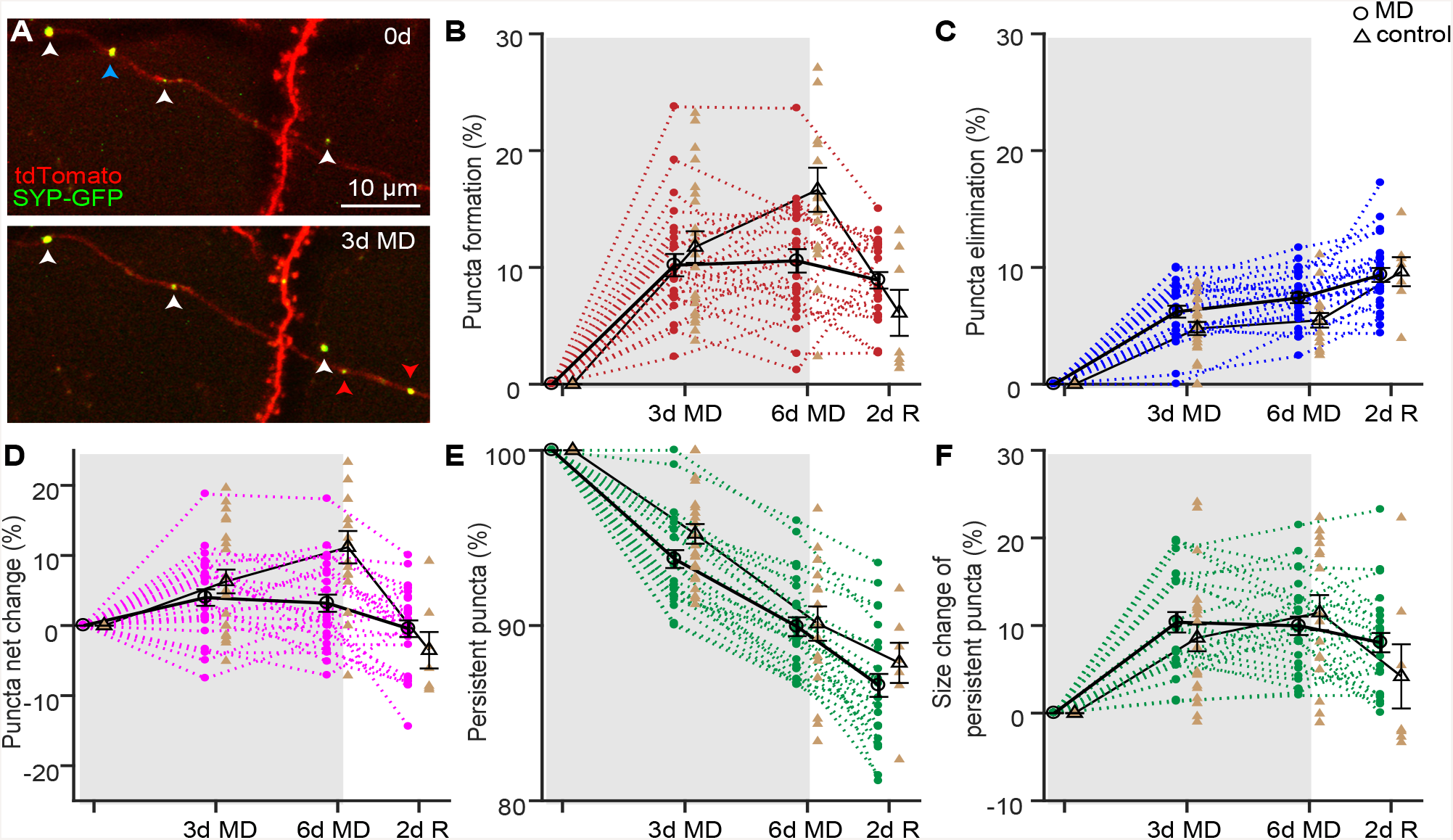
Changes in presynaptic boutons of layer 2/3 neurons. A, Presynaptic marker SYP-GFP co-expressed with tdTomato in a layer 2/3 excitatory neuron. White arrows mark persistent boutons, red arrows mark gain of boutons, and a blue arrow marks loss of a bouton. B-D, Percentage change in bouton formation, elimination, and total net change over 8 days in the binocular zone. E, Fraction of persistent SYP-GFP puncta throughout the period of MD and recovery. F, Percentage change in size of persistent SYP-GFP puncta. Each point marks the average value for the 17-61 bouton on a single axonal branch. Successive measurements of each branch in experimental animals connected by dotted lines. Average values for the 25 branches in 4 experimental animals indicated by open circles and heavy black line (864 boutons total). Control data plotted in light brown; average values for the 24 branches in 3 control animals indicated by open triangles connected by light black lines (727 boutons total).

In MD mice, 3 days of MD did not have a significant effect on bouton formation (10.2±1.1% in 3d MD vs 11.7±1.7% in 3d control, p = 0.34, n = 25 branches in 4 experimental animals, n = 24 branches in 3 control animals, unpaired Welch’s t-test) or elimination (6.2±0.6% in 3d MD vs 4.8±0.7% in 3d control, p = 0.064) compared to control. However, by 6 days of MD, bouton formation was significantly reduced from control (10.6±1.2% in 6d MD vs 16.6±2.5% in 6d control, p = 0.0035), and bouton elimination was increased (7.4±0.5% in 6d MD vs 5.5±0.8% in 6d control, p = 0.0014). Therefore, the net change in the numbers of presynaptic boutons was significantly affected by 6 days of MD (3.2±1.4% in 6d MD vs. 11.2±3.1% in 6d control, p = 0.0019) but not 3 days of MD (4.0±1.4 % in 3d MD vs. 6.3±2.1% in 3d control, p = 0.26). This temporally restricted increase is consistent with a strengthening of synapses and the homeostatic scaling mechanisms demonstrated to occur between 3 and 6 days of MD (16). Notably, MD did not have a significant effect on persistent boutons (Fig 5EF).

## Discussion

Previous physiological studies of ocular dominance plasticity have focused on the cortical neurons in the thalamorecipient layer 4 as well as the layers above and below (reviewed in (3, 6, 39)). However, studies of structural plasticity *in vivo* during the critical period have focused on the spines of apical dendrites in layer 1, in most cases from pyramidal neurons in layer 5 (37, 40). The physiological changes in these neurons were generally not characterized, making it difficult to determine the extent to which the time course of the structural changes was consistent with physiological plasticity and might therefore underlie it.

The present study examines both postsynaptic and presynaptic structures in axons and dendrites of layer 2/3 excitatory neurons, in which the time course of ocular dominance plasticity and underlying signaling mechanisms have been the subject of many reports (reviewed in (6)). To investigate the anatomical substrates that underlie the ocular dominance plasticity, we tracked the gains and losses of presynaptic boutons and postsynaptic densities in individual neurons in the binocular zone of mouse visual cortex during critical period. We labeled major proteins of dendritic spines (PSD95) and presynaptic boutons (SYP) in individual layer 2/3 neurons and followed them through 6 days of monocular deprivation during the peak of the critical period plus 2 days of recovery with binocular vision, the same deprivation and recovery protocol that was used for the studies of cellular signaling mechanisms (15, 16). To determine whether the anatomical changes corresponded with eye-specific visual responsiveness in individual neurons, we applied calcium imaging to measure the physiological effects of MD in neurons after measuring their anatomical changes. Our findings suggest that the rapid anatomical rewiring of intracortical connections serve as a structural basis for the physiological effects of monocular deprivation in these layer 2/3 neurons.

### Measuring structural changes in synapses below layer 1

Measuring changes in synaptic structures below the most superficial layer of the cortex using optical microscopy is challenging even with 2-photon microscopy. Because the best possible axial resolution is comparable to the size of the smaller dendritic spines, the presence of the fluorophore within both dendrite and spine may artifactually cause the spine to disappear or reappear when the size or orientation of a spine, or the clarity of the tissue, changes across imaging sessions. The use of fluorescent label for the major protein of the postsynaptic density (PSD95) produces discrete puncta in the image that can be counted unambiguously and measured accurately, independent of the inevitable changes in optical resolution within deeper cortical layers over days.

Overexpression of PSD95 is known to alter synaptic plasticity both in hippocampus and cerebral cortex (41–45). While our aim was to deliver only tracer amounts of fluorescent protein that would have no effect on plasticity, control experiments were required to exclude such an effect. We performed these by measuring cortical plasticity *in vivo* after delivering a much higher concentration of PSD95-GFP plasmid with much wider expression than was used in the actual experiments, and we found that the response to monocular deprivation was normal (Fig 1K).

The resolution of presynaptic boutons in optical microscopy is even more difficult than for dendritic spines because not all swellings of an axon are presynaptic sites. To count and measure presynaptic terminals we labeled synaptophysin, a major presynaptic protein essential for synaptic vesicle release (46–48). SYP-GFP appears as discrete puncta along axons in 2-photon microscopy (Fig 1, 5). As with PSD95-GFP, even much greater expression of SYP-GFP in a wider population than was used in the experimental cases did not affect normal ocular dominance plasticity (Fig 1M).

### Reorganization at layer 2/3 postsynaptic sites closely follows functional changes

Ocular dominance plasticity during early development takes places in three stages with distinctive physiological changes in layer 2/3 neurons in the binocular zone of V1 (Fig 6A): (1) a dramatic loss of response to the deprived eye, (2) a significant increase in open eye response and a slight increase in deprived eye response, and (3) a return of responses to their initial state after re-opening the deprived eye (reviewed in (6)).

**Figure 6.**
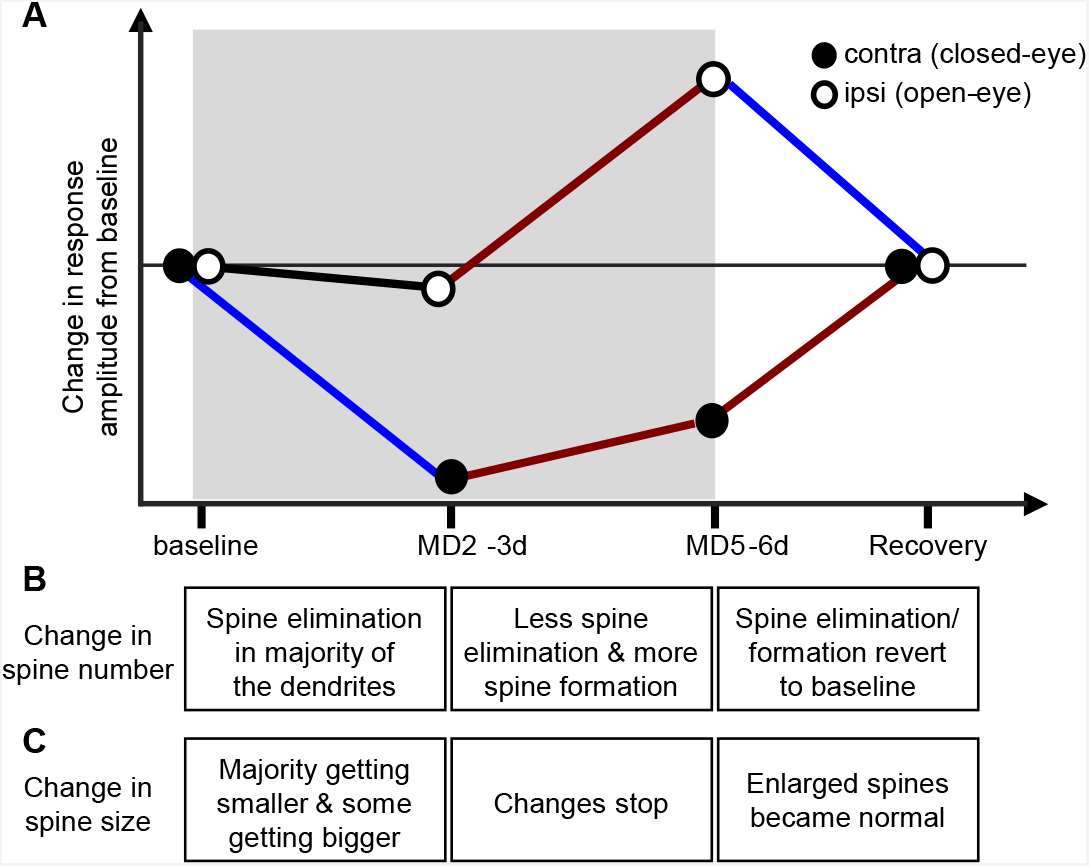
Stages of ocular dominance plasticity during critical period. A, Ocular dominance plasticity takes place in temporally and mechanistically distinct stages: a decreased response to the deprived eye in the first 2 to 3 days, a delayed increase in response to the open eye, and a full recovery in 2 days of normal vision. B, Change in spine number has comparable time course and direction: loss of spines by 3 days of MD, decreased spine elimination and increased spine formation by 6 days of MD, and a full recovery to baseline after 2 days of binocular vision. C, Size change in the of persistent spines reacts faster to MD: majority of dendrites has decreased spine size and some dendrites has increased spines sizes by 3 days of MD, and no further change by 6 days of MD, and enlarged spines become normal after restoration of normal vision.

The present study finds that structural changes in post-synaptic spines have a similar time course and direction of change to physiological ocular dominance plasticity (Fig 6BC). After 3 days of MD, there was a significant loss of spine number due to greater spine elimination. By 6 days of MD, there was more spine formation and less elimination, resulting in a small net change in spine numbers averaged over all the dendrites, but the changes in individual dendrites became so diverse that they could no longer be fit by a normal distribution. The sizes of persistent puncta also became very diverse at both 3 and 6 days of MD, suggesting that enlargement of some of the existing puncta precedes the formation of new spines in response to altered visual inputs. Restoration of binocular vision for 2 days quickly brought back these diverse changes to a normal distribution in size and number.

The neurons in which calcium imaging of visual responses was carried out following structural measurement demonstrated a strong correlation between each neuron’s spine remodeling and physiological responses to the two eyes. Previous studies have demonstrated that spines from the same dendritic tree tend to share the same orientation preference, while different dendritic segment from the same neuron are biased toward different orientations (49). If inputs to individual dendrites were also eye-specific, then MD would be expected to produce diverse changes in spine number and size on different dendrites, as we observed (Fig. 3D). The gains and losses of PSD95-GFP puncta in individual dendrites of deprived animals showed a tendency toward anticorrelation that did not reach statistical significance. Our functional imaging using OGB1-AM, while revealing somatic responses, was not capable of resolving dendritic signals. It will be interesting for future studies to explore potential dendritic specificity for ocular dominance.

Our findings of changes in spines of individual neurons are consistent with population studies of developing visual cortex showing decreased spine density after visual deprivation by MD (36, 37) or dark-rearing (34, 35). A number of studies (33, 35, 36) have shown correlations between anatomical and functional changes at the population level consistent with our demonstration of a relationship between anatomical change and eye-specific responses at single cells. The rapid remodeling, including changes in number and size of spines, and the correlation between structural and functional changes in individual cells observed in the present study suggest that these changes may be the cellular substrate for functional reorganization during experience-dependent plasticity in the developing mouse V1 (Fig 6).

### Delayed reorganization of layer 2/3 presynaptic boutons during the critical period

Control mice receiving normal visual inputs showed a steady increase of presynaptic bouton number during the critical period, whereas the number of postsynaptic spines in control animals remained stable during this period. It is worth noting that, while we specifically labeled presynaptic and postsynaptic structures in layer 2/3 excitatory neurons, the synaptic markers used, SYP-GFP and PSD95-GFP, do not target the same types of synapses: PSD95-GFP puncta receive excitatory inputs, principally from layer 2/3 and layer 4 excitatory neurons, though these spines may also receive some degree of inhibition. On the other hand, the presynaptic structures labeled by SYP-GFP target inhibitory as well as excitatory neurons, both within layer 2/3 and possibly onto the dendrites of neurons in other layers. Structural changes in the presynaptic outputs from layer 2/3 cells, unlike those in their postsynaptic inputs, might therefore not be expected to match the changes in their visual responses. These differences may explain why the effect of MD on the presynaptic boutons in the cells examined was smaller and slower than the changes at postsynaptic spines.

Many studies have shown that maturation of inhibitory neurons at the end of critical period is involved in critical period plasticity (3, 39, 50). The increase in bouton formation at the end of the critical period when spine number on the excitatory neurons remains stable suggests an increase of excitatory inputs onto local inhibitory neurons, which may contribute to the end of the critical period. In the adult visual cortex, 2-photon microscopy of gephyrin-labeled inhibitory synapses show them to be highly dynamic, being removed and replaced repeatedly at the same sites, in contrast to the relatively stable excitatory synapses (32). Adult ocular dominance plasticity is associated with a specific loss of inhibitory inputs (29, 30). It will be interesting to apply similar techniques to measure the turnover of inhibitory synapses during the critical period.

## Methods

### Experimental animals

All animal work was approved by the University of California San Francisco Animal Care and Use Program and conforms to the National Institutes of Health guidelines. Ai14 (Stock No: 007908, Jackson Laboratory), and wild-type C57BL/6J (Charles River, CA) lines were purchased to breed and maintain in UCSF housing facility, on a standard 12h dark/ light cycle. Both sexes of animals were included in the study.

### DNA constructs and *in utero* electroporation

The three plasmids used in our present study, pCAG-Cre, pEF1a-double flox-PSD95-eGFP-WPRE, and pCAG-double floxed-SYP-eGFP-WPRE were generated by subcloning original constructs and then customized into specific promoters (available in Addgene for distribution). Plasmids were delivered through *in utero* electroporation as previously described (51, 52) to E16 Ai14 mouse embryos to label layer 2/3 cortical excitatory neurons. Pregnant female mice were anesthetized with isoflurane (3% for induction and 1.5% for maintenance) with injection of Buprenex (0.1 mg./kg.) and Carprofen (15 mg./kg.). After laparotomy, approximately 1μl of a solution consisting of ACSF, 0.1% of Fast Green (Sigma-Aldrich, St. Louis, MO, USA), pCAG-Cre (0.0001 μg/μl), and one type of double flox marker (0.75 μg/μl) was injected into the left ventricle of each embryo through a glass capillary with a tip of diameter 20-30 μm (Drummond Scientific, Broomall, PA). A pair of platinum electrodes positioned to target the dorsolateral wall of the left hemisphere and a series of five square-wave current pulses (35 V, 50 ms duration, 950ms interval) were delivered by a pulse generator (ECM830; BTX, San Diego, CA). Afterwards, embryos were put back to the mother’s womb and sutured up for normal delivery and growth.

### Headplate and cranial window implantation

For long-term visualization of neuronal morphology under 2-photon microscope, animals around P21 (4-5 days before imaging) were implanted with a stainless headplate and a cranial window was made as previously described (53–55). Briefly, animals were anesthetized with isoflurane (3% induction; 1.5% surgery), and injected subcutaneously with atropine (0.15 mg./kg.), dexamethasone (1.5 mg./kg.), and carprofen (15 mg./kg.). After a scalp incision and fascia cleaning, customized stainless steel headplate was secured to the skull with cyanoacrylate glue (VetBond, 3M) and dental acrylic (Lang Dental Black Ortho-Jet powder and liquid). A craniotomy was made with steel drill bits centering approximately on the binocular zone of the visual cortex (3-mm lateral to midline, 1mm anterior to lambda), gel foam soaked in ACSF was used to cover the exposed cortical area for 5 min, and a 3-mm diameter circular glass cover slip was secured using crazy glue.

### Intrinsic signal imaging

Binocular V1 was functionally located using a brief transcranial intrinsic signal imaging as previously described (15, 56). To evoke robust visual responses, mice were injected with chlorprothixene (2 mg/kg, i.m.) and maintained anesthesia using a low concentration of isoflurane (0.6%–0.8% in oxygen), the core body temperature was maintained at 37.5° using a feedback heating system. A temporally periodic moving 2°-wide bars was generated using the Psychophysics Toolbox (57, 58) in Matlab (Mathworks) and continuously presented at a speed of 10°/sec. To identify binocular V1, the visual stimulus subtended 20° horizontal and was presented to one eye at a time (the non-tested eye blocked with eye shutter) with monitor positioned directly in front of the animal. To calculate ODI, response amplitude was averaged from at least four measurements, and ODI was computed as (R − L)/(R + L), where R and L was the peak response amplitudes through the right eye and the left eye, respectively, using a custom Matlab code (59).

### Monocular deprivation

Monocular deprivation was induced by suturing the right eyelid with 7-0 polypropylene monofilament (Ethicon). Suture integrity was inspected immediately prior to each imaging session. Animals whose eyelids did not seal fully shut or had accidentally reopened were excluded from further experiments.

### Two-photon imaging

Two-photon imaging was performed using a Movable Objective Microscope manufactured by the Sutter Instrument Company. A mode-locked Ti:Sapphire laser (Chameleon Ultra 2, Coherent Inc.) was tuned to 910nm, and the laser power through the objective was adjusted within the range of 50–100 mW. Z-resolution was controled with a piezo actuator positioning system (Piezosystem Jena, Germany) mounted to the objective. Emission light was collected by a 40X water-immersion objective (NA0.8; IR2; Olympus), filtered by emission filters (525/70 and 610/75 nm; Chroma Technology) and measured by two independent photomultiplier tubes (Hamamatsu R3896). Scanning and image acquisition was controlled by ScanImage software (Vidrio Technologies). Mice were anesthetized with isoflurane and head-fixed for structural and functional imaging.

### Imaging processing pipeline and semi-automated puncta detection

Segmented images from the same imaging sessions were first stitched using linear transformation and then images from different imaging sessions were further registered with rigid alignment and 3-dimensional warping (38). To compare GFP-tagged synapses across imaging sessions, a customized and semi-automated ImageJ plugin (deposited at https://github.com/ucsfidl/Puncta_Counter) was used to outline and group the 2-dimensional boundaries of individual synapses to create 3-dimensional boundaries. Identified synapses had to be a minimal size of 2×2×2 or 8 clustered pixels with a minimal average signal intensity of at least three times above shot noise background levels, consistent with synapses of these pixel dimensions correlating to their true physical dimensions as measured after EM reconstruction (29). Using these criteria, the software automatically identifies and outlines at least 95% of synapses, with almost no false positive identification of background noise or ascending neurite branch. Less than 1% of two or more synapses are interpreted as one synapse and less than 1% of single synapses are interpreted as two synapses.

Individual synapses are given a unique identification number and a number corresponding to cell of origin identified by using tdTomato neurite labeling. Following the identification and outlining of synapses, the position, pixel volume (total number of pixels), and GFP intensity (sum of all pixel values within outlined synapse following application of a 0.58 **μm Gaussian blur) were calculated. The GFP inten**sity was normalized to the average tdTomato intensity of all planes that include the synapse in order to compensate for differences in optical clarity and laser power during imaging. Stable synapses were defined as synapses that were identified in all imaging sessions. For stable synapses making projections to a single cell, the pixel volume and the normalized GFP intensity were compared across imaging sessions and used to determine changes in synaptic strength. Dynamic synapses were defined as synapses that appeared or disappeared for one or more imaging sessions.

Axonal or basal dendritic branches containing at least 15 PSD95-GFP or SYP-GFP puncta within a 40 μm range of depth were selected for analysis. This plane included the soma in the case of the dendrites. Changes in puncta numbers were measured from the baseline images. The net change was calculated as the numbers of puncta added minus those eliminated divided by the baseline numbers. Puncta were counted as persistent if they were continuously present from the baseline images.

### Calcium imaging and data analysis

In a separate group of animals, we also measured the visual responses of the neurons that underwent structural imaging to correlate the functional and anatomical relationship. To obtain functional responses of the labeled neurons, calcium-sensitive dye Oregon Green Bapta1-AM (OGB1-AM; Molecular Probes) was applied after structural imaging. Glass coverslip over the cranial window was carefully removed with fine forceps after drilling off dental acrylic. As previously described (60), OGB1-AM (dissolved in DMSO containing 20% Pluronic F-127 and further diluted 1:10 in bath solution 50 NaCl, 2.5 KCl, 10 Hepes, pH 7.4) was slowly injected into binocular zone under constant pressure of ~10 psi (Picospritzer, General Valve) for ~1 min at a depth of 150-300 μm through a pipette with a tip size of ~2 μm (inside diameter). The progress of administration was visualized by inclusion of a nontoxic red fluorescence dye, Alexa 594 (Invitrogen). Typically, three injections were made to cover the binocular zone.

To measure the eye-specific responses under two-photon microscope, 2.5% agarose in ACSF was applied over the brain, sealed with a coverslip and petroleum jelly. A preanesthetic injection of chlorprothixene (2 mg. /kg., i.m.) and 0.7% isoflurane was applied to allow robust visual responses. Full-field drifting sinusoidal gratings (0.04 cpd, 99% contrast, 1 Hz temporal frequency) at 8 evenly spaced directions were generated and presented for 3 sec, separated by 3-sec blank periods of uniform 50% gray, in random sequence using the MATLAB Psychophysics Toolbox (57, 58). Stimuli to the two eyes were randomly interleaved using a pneumatic shutter and the stimulus sequence was presented 5 times, randomizing the prodder between presentations. Independent component analysis was then used to extract somatic calcium transients as a measure for cellular visual responses (61)with ensembles of cells arranged from the surface to the white matter. Within a column, neurons often share functional properties, such as selectivity for stimulus orientation; columns with distinct properties, such as different preferred orientations, tile the cortical surface in orderly patterns. This functional architecture was discovered with the relatively sparse sampling of microelectrode recordings. Optical imaging of membrane voltage or metabolic activity elucidated the overall geometry of functional maps, but is averaged over many cells (resolution >100 microm. To calculate ODI, response amplitude was averaged from at least four measurements, and ODI was computed as (R_p_ − L_p_)/(R_p_ + L_p_), where R_p_ and L_p_ were the peak response amplitudes at the preferred orientation through the right eye and the left eye, respectively.

### Statistical Analysis

Distribution of group data was tested for normality using Lilliefors test. Only data that passed normality test were presented as mean ± SEM. For comparison between experimental and control groups, unpaired Welch’s (unequal variance) t-test was conducted with p value reported. For comparison of change in the same branches between experimental and control groups over days, unbalanced 2-way ANOVA was conducted, followed by post hoc tests using Tukey’s method, and the adjusted p value was reported. For function and structure relationship, regression was performed using Matlab linear least squares fitting, with adjusted R-squared reported.

## Acknowledgements

We thank Drs. K. Svoboda (PSD95), S. Arber (Synaptophysin), and K. Deisseroth (Double flox) for the original constructs and R. Mostany, C. Portera-Cailliau, and V. Cairone for technical consultation. We thank members of the Stryker laboratory for helpful discussion, particularly M. Kaneko, S. Gandhi, and C. Niell for assistance and training on *in utero* electroporation and calcium imaging. Supported by grants from the U.S. National Institutes of Health R01EY02874 (to M.P.S.), K22NS089799 (to J.S.E.), and K99EY029002 (to Y.S.).

## Notes

#### Summary of Updates

Clarification of methods, statistics, and introduction. Correction of error in figure caption.

https://github.com/ucsfidl/Puncta_Counter

